# Environment by environment interactions (ExE) differ across genetic backgrounds (ExExG)

**DOI:** 10.1101/2024.05.08.593194

**Authors:** Kara Schmidlin, C. Brandon Ogbunugafor, Alexander Sastokas, Kerry Geiler-Samerotte

**Affiliations:** Biodesign Center for Mechanisms of Evolution, Arizona State University, Tempe, AZ, 85287; School of Life Sciences, Arizona State University, Tempe AZ, 85287; Department of Ecology & Evolutionary Biology, Yale University, New Haven, CT,06511; Santa Fe Institute, Santa Fe, NM, 87501

## Abstract

While the terms “gene-by-gene interaction” (GxG) and “gene-by-environment interaction” (GxE) are widely recognized in the fields of quantitative and evolutionary genetics, “environment-byenvironment interaction” (ExE) is a term used less often. In this study, we find that environmentby-environment interactions are a meaningful driver of phenotypes, and moreover, that they differ across different genotypes (suggestive of ExExG). To support this conclusion, we analyzed a large dataset of roughly 1,000 mutant yeast strains with varying degrees of resistance to different antifungal drugs. Our findings reveal that the effectiveness of a drug combination, relative to single drugs, often differs across drug resistant mutants. Remarkably, even mutants that differ by only a single nucleotide change can have dramatically different drug x drug (ExE) interactions. We also introduce a new framework that more accurately predicts the direction and magnitude of ExE interactions for some mutants. Understanding how ExE interactions change across genotypes (ExExG) is crucial not only for modeling the evolution of pathogenic microbes, but also for enhancing our knowledge of the underlying cell biology and the sources of phenotypic variance within populations. While the significance of ExExG interactions has been overlooked in evolutionary and population genetics, these fields and others stand to benefit from understanding how these interactions shape the complex behavior of living systems.

## Introduction

Over 100 years ago, William Bateson (1) used the term, “epistasis,” to describe peculiar findings where the phenotypes of offspring deviated from expectation in a way that could not be accounted for by dominance effects nor differences in environment (2). More recently, the term “epistasis” has come to include any genetic interaction (GxG) where the combined effect of two genetic changes differs from the sum of their individual contribution (2, 3). Or, as one colloquial definition frames it, epistasis is the “surprise at the phenotype when mutations are combined, given the constituent mutations’ individual effects” (4). Genetic interactions have been of interest, in both classical and modern settings, because they complicate a major goal of biology: predicting phenotype from genotype (5–8). Scientists have debated the impact of genetic interactions on such prediction efforts (9, 10) and which types of interactions, e.g. gene x gene (GxG) or gene x environment (GxE), are important (11). These interactions are of interest to other disciplines as well (12). For example, genetic interactions have suggested which genes participate in the same regulatory modules (13, 14), predicted which evolutionary trajectories are most likely (3, 15), and revealed global constraints on protein evolution (16) and adaptive evolution (17). Given their broad utility to biologists, many useful mathematical frameworks exist for quantifying GxG (18), GxE (19) and GxGxE (3, 11, 20). Further, many experimental frameworks have comprehensively surveyed GxG or GxGxG (15, 16, 21–23), GxE (24–27), or GxGxE (24, 28–31). But one type of interaction has remained largely neglected by quantitative geneticists: ExE interactions, or those arising from interactions between environments (**Figure 1A**).

**Figure 1:**
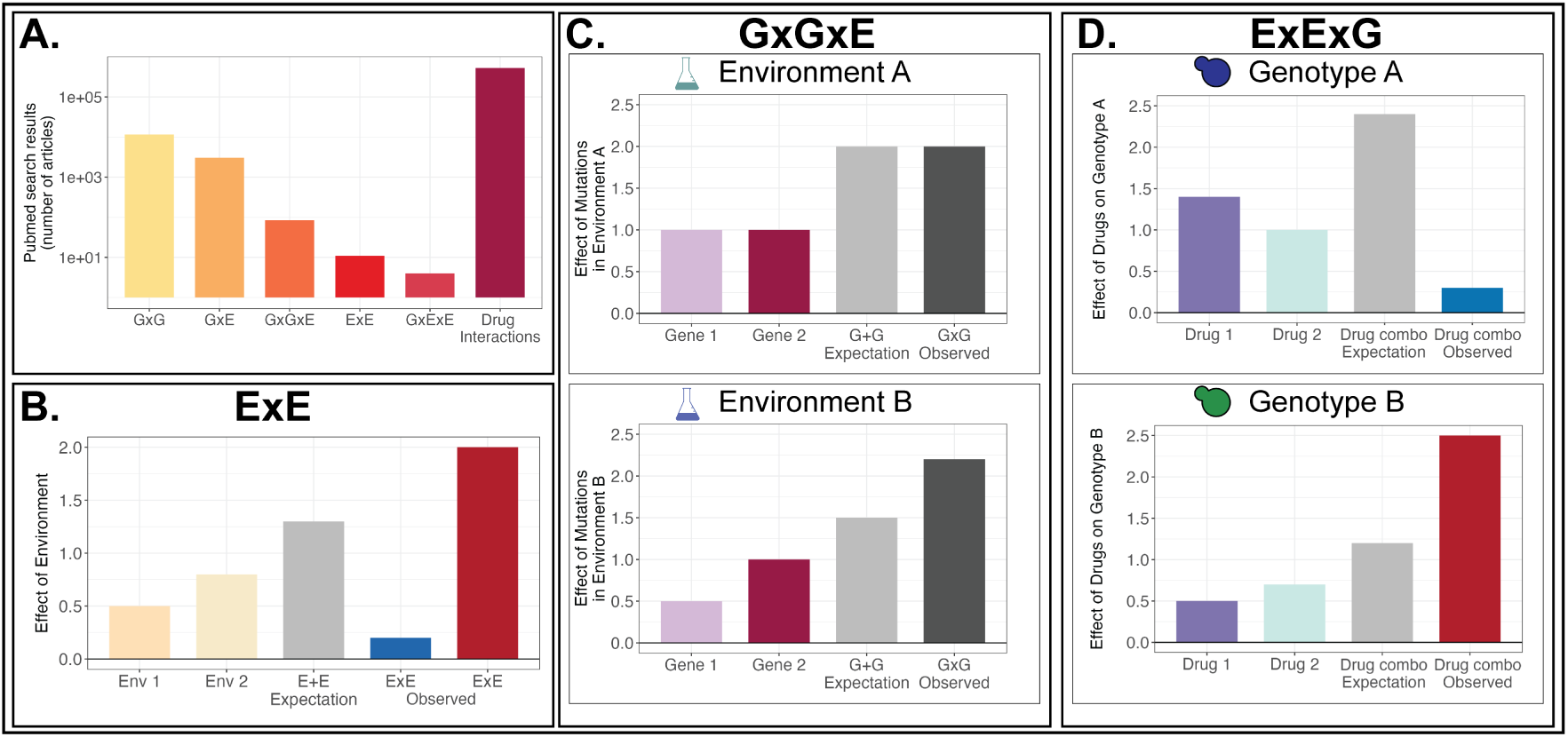
Comparative visualizations of ExE, GxGxE and ExExG interactions. **(A)** ExE interactions are understudied. Search results retrieved from Pubmed on May 3, 2024 demonstrate that publications describing ExE interactions, including GxExE, show substantial disparities when compared to simpler interactions like GxG and GxE, and drug interactions, which have significantly greater representation. Complete search term results are located in table S1. **(B)** A cartoon to define ExE. Environments 1 and 2 have unique effects on an organism’s phenotype or fitness (light orange and light yellow bars). When exposed to both environments simultaneously, one might expect that the combined effect is additive (E+E, indicated by gray). Here, we define ExE as when the observed effect of combining environments differs from the expectation (blue and red bars). **(C)** A cartoon to define GxGxE. GxGxE interactions describe how the combined effect of the same two mutations (light pink and dark pink bars) changes across two or more environments (top vs bottom panels). In this cartoon, the effects of gene 1 and gene 2 are additive in environment A (top panel; expectation equals observed), but produce unexpected interactions in environment B. Since the interaction between genes (GxG) differs across environments, this is referred to as a GxGxE interaction. **(D)** A cartoon to define ExExG. In general, ExExG interactions describe how the combined effect of two environments (purple and teal bars) changes across two or more genetic backgrounds (top vs. bottom panels). In this manuscript, the environments we study are different drugs. Different drug-resistant genotypes are exposed to the same single drugs (Drug 1, purple and Drug 2, teal) and their combination (Drug combo, gray). In this cartoon, genotype A (top) is resistant to drug 1 and 2 and thus has a fitness advantage over the ancestor of all the drug-resistant mutants in these environments (purple and teal bars). But genotype A is unexpectedly sensitive to the combination of these two drugs, losing almost all of its fitness advantage (blue bar). This might imply that Drug 1 and drug 2 interact synergistically, enhancing one another’s ability to harm cells. However, this is not the case for genotype B, with respect to which the drugs interact antagonistically, meaning they hinder one another’s ability to harm cells, resulting in genotype B having an increased fitness advantage over the ancestor (red bar). Since the effect of combining drugs (ExE) varies across genotypes, this is referred to as ExExG.

Here, we define ExE (i.e. environment-by-environment interactions) as when the combined effect of two environments on phenotype is unexpected given their individual effects (**Figure 1B**). For example, if a microbe grows slowly in a high salt environment and equally slowly in a high temperature environment, but does not grow even slower in a high salt plus high temperature environment, this would be unexpected under an additive model and herein termed “ExE”. Perhaps the reason for the near omission of the term “ExE” in the quantitative genetics literature is straight-forward: there is no genetic component (no “G”), so those who map the effects of genetic changes onto phenotype are naive (or disinterested) to the benefits of quantifying ExE interactions. But there are several reasons it may be worthwhile to turn attention towards ExE. For one, understanding why environments have non-additive effects on phenotype stands to expand knowledge about regulatory network architecture (32, 33), as have GxG and GxE models (13, 34). Further, if ExE often varies across genetic backgrounds, in other words, if ExExG is common, then quantitative and evolutionary geneticists can incorporate ExExG interactions into models that predict the phenotypic effects of mutation. ExExG is not the same phenomenon as GxGxE (**Figure 1C–D**). Several studies have examined the power of GxGxE interactions, or the role of the environment in sculpting epistatic interactions (labeled “environmental epistasis”; see Lindsey et al 2014) (11, 24, 30, 35). To date, only a handful of studies mention ExExG (36–42), though usually not in a way that speaks to the circumstance whereby different genotypes tune the interactions between environments (the focus of the current study).

One key reason to study ExE pertains to understanding how multidrug environments affect microbial phenotypes (43–45), though in the relevant literature ExE interactions are usually termed “drug interactions” (32, 46) or occasionally “drug epistasis” (47) rather than “ExE” (**Figure 1A**). There is practical interest in finding pairs of drugs that interact ‘synergistically’, i.e., the combination of both drugs is more effective than one would predict based on either single drug (**Figure 1D**; top panel) (48–52). But just as genotype-phenotype mapping studies rarely examine environment interactions, drug synergy studies focus on genetic interactions less frequently. For example, several studies suggest that if one understands the cell biological mechanisms underlying drug interactions, one can predict synergy (53–55), but this ignores that mutations may change the underlying drug interactions (56, 57). Studies of the combined impacts of multiple environmental stressors on natural ecosystems often make a similar omission (58, 59). Other studies describe the biggest challenge in detecting synergy as there being more possible combinations of environments than one can study (44, 53, 60), but this ignores that studying these combinations in multiple genetic backgrounds would be even more difficult. Despite the combinatorics challenge, efforts have been made to measure large numbers of drug and environment interactions (58, 60), including higher-order interactions (61, 62), which have fueled multidrug treatment strategies and evolutionary models (63). But these treatments and models could fail if mutations change the way environments interact (57).

Indeed, the literature describes several cases where drugs interact differently across different mutants or cell lines (60). For example, the antifungal drugs fluconazole and radicicol each administered independently have little effect on the fitness of erg3 mutants in yeast, but act synergistically to kill these mutants (64, 65). However, numerous yeast strains resist fluconazole via mutations such as those to PDR3 and ERG11 that are less sensitive to the addition of radicicol (56, 66). Similarly, recent screens for other types of drug interactions, e.g., collateral sensitivity, have shown that these interactions can change dramatically across different drug-resistant mutants (67). Further study of the extent to which drug and environmental interactions change across genetic backgrounds (ExExG), and deeper consideration of how this affects predictive models, is needed.

Large-scale study of ExExG has recently become possible due to evolution experiments that utilize DNA barcodes (56, 68) to create thousands of adaptive microbial strains that each possess only a small number of genetic differences and are highly tractable, meaning their fitness relative to a common ancestor can be measured in many conditions using pooled barcoded competitions. Here, we take a large collection of roughly 1,000 antifungal drug resistant yeast mutants evolved using this method and ask how often fitness in multidrug environments is predicted by fitness in single drug environments (**Figure 1D**). We find substantial ExE (i.e., multidrug fitness is not easily predicted by single drug fitness). We also find substantial ExExG (i.e. the magnitude and direction of ExE are different across different mutants). We demonstrate that single point mutations often alter ExE and that even similar adaptive mutants that emerge from the same evolution experiment can have different ExE.

Given the prevalence of ExExG in our data, we next explored some new ways to study ExE and ExExG. We applied a GxG model to better predict environmental interactions for some mutants. We also observed that diverse mutants cluster into groups with similar ExE, implying the ExE of some mutants can be used to predict ExE of others. In general, our findings call for greater study of ExExG across disciplines, including among scientists interested in modeling the evolution of drug resistance, the links from genotype to phenotype (5), how gene expression responds to environmental change (36), the construction of microbial communities (69), and how the interaction between different forces crafts complex biological systems (6).

## Results

### Environment by environment (ExE) interactions vary across drug pairs

In order to study environment-by-environment interactions, we compared data from pooled fitness competitions conducted in 4 environments each containing a single drug to data from 4 environments representing all pairwise combinations of these drugs (56) (**Figure 2A**). We asked if multidrug fitness of 1000 drug-resistant mutants was easily predicted by fitness in each single drug environment. We used four different models (**Figure 2B**) to predict fitness in the drug combination environments, including the simple additive model depicted in **Figure 1** and other common models (32, 43, 44, 52, 70). None of the models we tried accurately predicts fitness in all four drug combinations. For example, fitness in the combined low rad + low flu environment (LRLF) is often predicted by taking the higher fitness of the low rad and low flu single drug environments (**Figure 2B**; leftmost panel; median falls on the zero line when using the highest single agent “HSA” model). But this same model tends to overpredict fitness in the high rad + low flu environment and underpredict fitness in the low flu + high rad environment (**Figure 2B**; middle panels; medians of HSA model fall farther from the zero line). Overall, there appears to be a good deal of ExE interaction. In other words, there are many cases where fitness in multidrug environments is not predicted by fitness in single drug environments.

**Figure 2:**
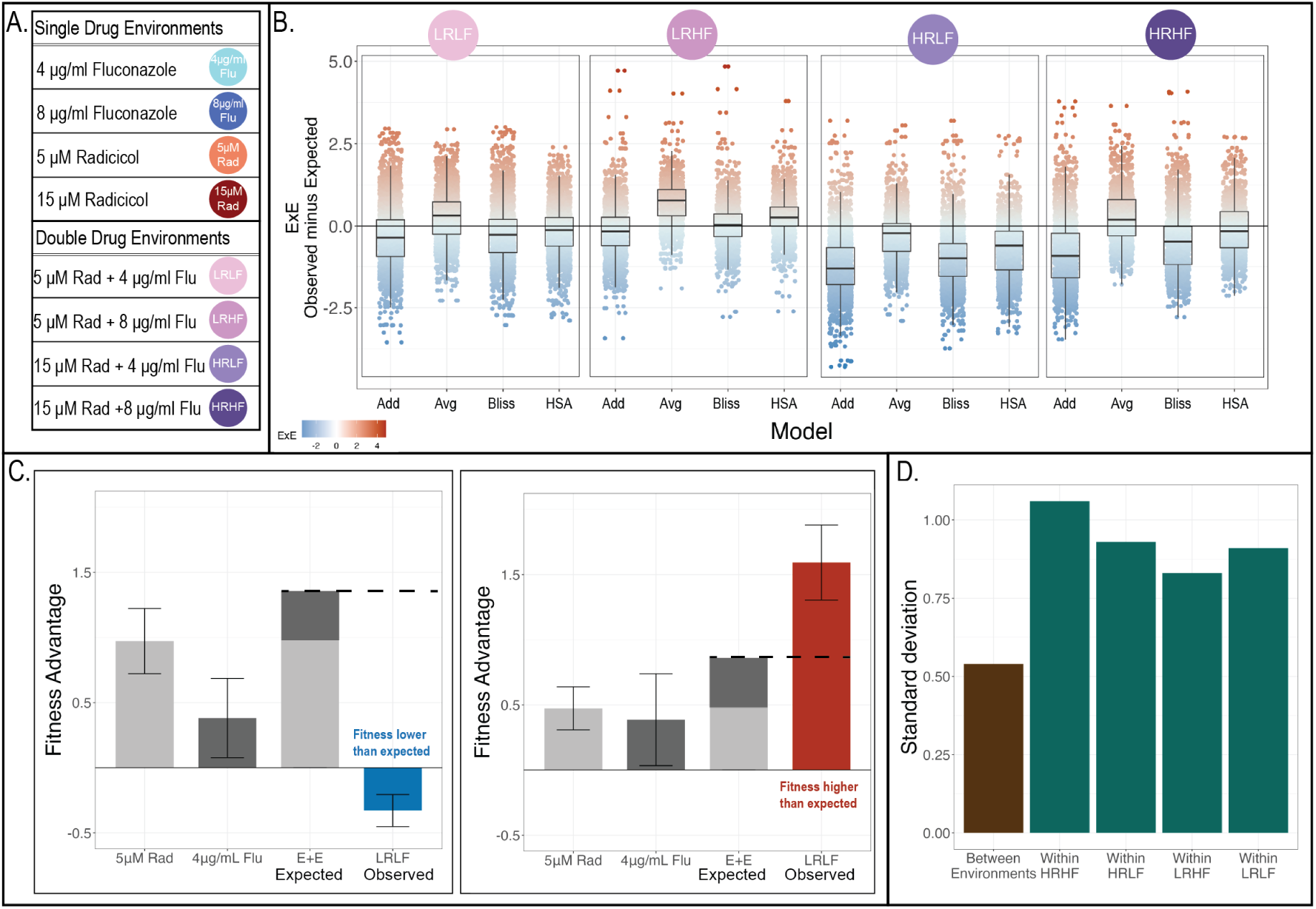
ExE interactions vary across drug pairs and across mutants. **(A)** We predict fitness in four double drug environments from fitness in four relevant single drug environments. **(B)** Environment-byenvironment interactions are revealed when fitness in a double drug environment deviates from the expectation generated by the relevant single drug environments. Four different models (horizontal axis) are used to calculate expected fitness for each of roughly 1000 mutants per drug pair (LRLF: n=1688; LRHF: n=850; HRLF: n=1318; HRHF: n=1023). Points representing each mutant are colored blue when a mutant’s fitness is worse than expected (synergy), and red when fitness is higher than expected (antagonism). Boxplots summarize the distribution across all mutants, displaying the median (center line), interquartile range (IQR) (upper and lower hinges), and highest value within 1.5 × IQR (whiskers). **(C)** Some mutants have different ExE interactions than others. The left panel displays the fitness of a yeast strain with a mutation in the HDA1 gene. It has lower fitness in the LRLF double drug environment than expected based on the simple additive has higher fitness than expected in the same environment. Error bars represent the range of fitness measured across two replicate experiments. Fitness is always measured relative to a reference strain, which is the shared ancestor of all mutant strains. **(D)** ExE interactions vary more across mutants than they do across drug pairs. The vertical axis displays the standard deviation across all four environments (brown) or across all roughly 1,000 mutants (green) when ExE is predicted using an additive model.

Like previous studies, we noticed that the direction of ExE interaction is sometimes specific to a multidrug environment (33, 61). For example, most of the models we tried tend to overpredict fitness in the high rad + low flu environment (HRLF). In other words, this combination of drugs is “synergistic”, meaning it hinders fitness more than expected based on the fitness effects of both single drugs (**Figure 2B**; third panel, more points are blue and most boxplot medians fall below the zero line). The opposite tendency, “antagonism”, appears more common in the low rad + high flu environment (LRHF). Fitness in this drug combination is often greater than expected based on fitness in the relevant single drug conditions (**Figure 2B**; second panel, more points are red and more boxplot medians fall above the center line). These trends are important because identifying synergistic drug combinations (those that are more detrimental than expected) could be helpful in treating viral (71), bacterial (72), and fungal infections (73), and cancers (60). Identifying drug pairs that interact antagonistically could be helpful as well by suggesting functional relationships between drug targets and strategies for restraining the evolution of drug resistance (32, 33, 61).

But, the major question of this study is: to what extent is synergy or antagonism a property of a drug pair? Even for drug pairs in which most of the mutants we study have lower fitness than expected, there are a few mutants that have unexpectedly high fitness (**Figure 2B**; there are always a number of red points even when most points are blue). So we next asked to what extent ExE varies across drug pairs versus across different mutants.

One additional thing to note from **figure 2B** is that different models often make generally different predictions. For example, the simple additive model tends to overpredict fitness in all four environments (**Figure 2B**; boxplots labeled “Add” are under the zero line). However, the average model is more likely to underpredict fitness (**Figure 2B**; boxplots labeled “Avg” are often above the zero line). Many previous studies discuss the strengths and weaknesses of these different models (52, 70), therefore, we do not focus on comparing models in this study. Our main focus here is that, no matter which model we use, we see mutants that deviate from the prediction in both directions, suggesting the presence of ExExG.

### ExE interactions vary more across mutants than they do across drug pairs

The drug resistant mutants we study were created in previous work by evolving a barcoded ancestral yeast strain in 12 different environments, including the 8 in **figure 2A** (56). Each mutant yeast strain differs from their shared ancestor by, on average, a single point mutation (56, 68). Yet, despite this similarity at the genetic level, there is variation in ExE (**Figure 2B**; see spread of points along vertical axis). To point to an example, one of these evolved yeast strains has a single point mutation in the HDA1 gene. It has unexpectedly low fitness in the LRLF environment given its fitness advantage in the relevant single drug environments (low rad: 5uML Rad and low flu: 4ug/mL Flu) (**Figure 2C**; left panel; error bars reflect range across 2 replicates). However, another (unsequenced) one of these evolved mutants has unexpectedly high fitness in this environment (**Figure 2C**; right panel; error bars reflect range across 2 replicates). The fitness of all mutants is measured relative to a reference strain, which is their shared ancestor (56).

While our previous work focused on 774 mutants with high quality fitness measurements in all 12 environments, here we are able to expand that collection. We do so by allowing each drug pair to have a unique dataset consisting of all mutant strains for which fitness was robustly measured in the relevant double and single drug conditions, plus a control condition with no drugs (LRLF: n=1688; LRHF: n=850; HRLF: n=1318; HRHF: n=1023). These datasets include 810 overlapping mutants for each of which we calculated ExE in all four drug pairs.

Overall, we found that ExE interactions vary at least as much across genotypes as they do across drug pairs. When using a simple additive model, the median amount of ExE varies across environments from -1.35 in HRLF to -0.3 in LRHF, with a standard deviation across all 4 drug pairs of 0.52 (**Figure 2D**; leftmost bar). This standard deviation is smaller than the standard deviation across mutants within each environment, which ranges from 0.8 to 1.05 (**Figure 2D**). In sum, these results suggest that ExExG is prevalent. Our follow-up analyses provide additional evidence that ExExG indeed reflects how ExE varies across different genes and strains.

### Mutations in different genes have different ExE interactions

Of the 810 drug resistant yeast strains present across all environments we survey, 53 have been previously sequenced at high enough coverage to identify the single nucleotide mutations that likely underlie drug resistance (56). A few genes appear to be common targets of adaptive mutation such that we can ask whether mutants in the same gene tend to have similar ExE interactions. For example, 35/53 sequenced drug-resistant strains have different mutations to either the PDR1 or PDR3 paralogs.

Other genes, such as SUR1, GBP2 and IRA1, were also found to be mutated in multiple different strains, though far less frequently than PDR1/3. Mutations to the same gene tend to have similar effects on fitness (**Figure 3 A–D**; error bars reflect standard deviation across all strains with mutations to a given gene).

**Figure 3:**
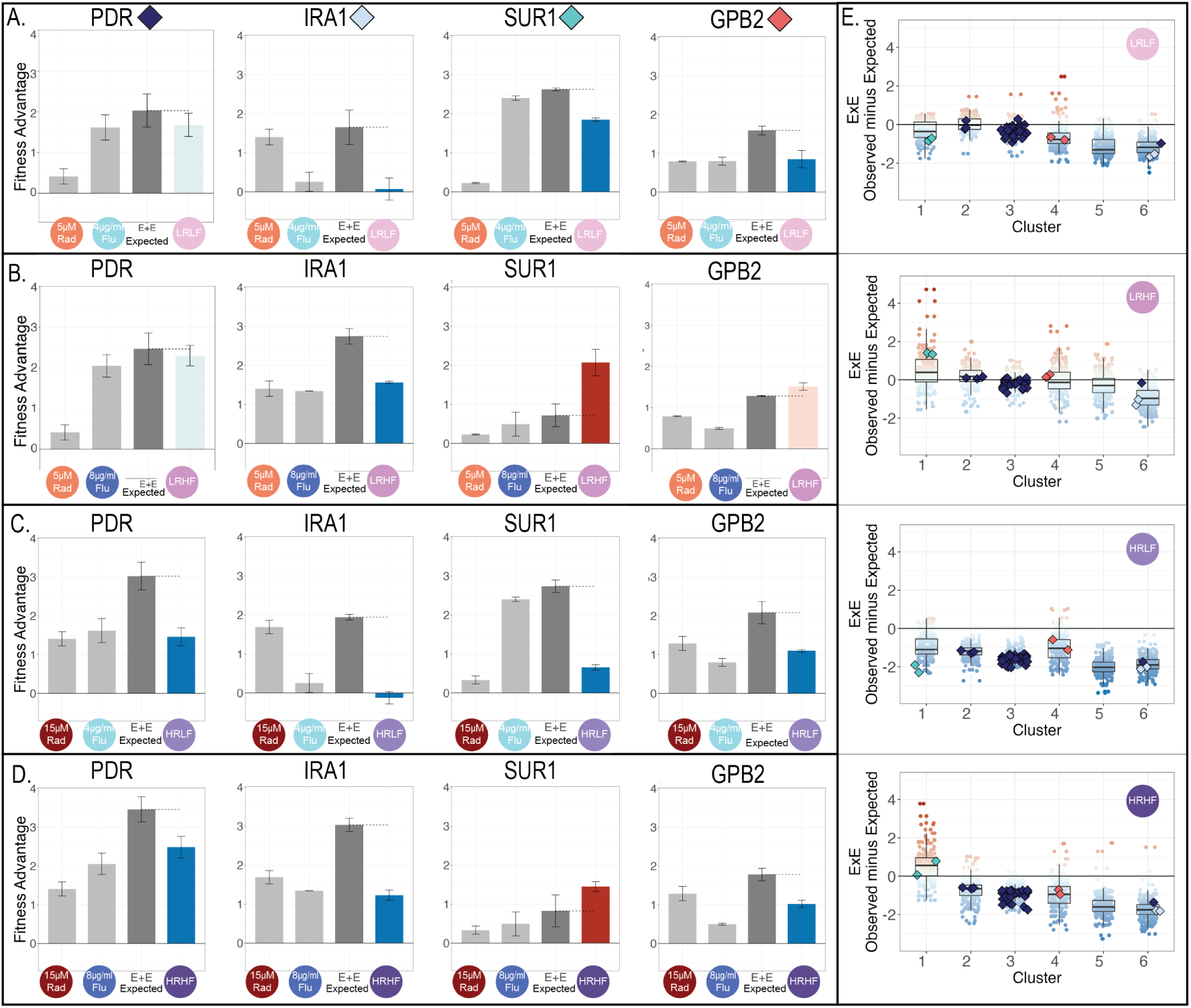
A few mutations can change a drug pair from having a synergistic to an antagonistic effect. (A –. **D)** Fitness advantages of strains with mutations in either PDR1/3 (n=35), IRA1 (n=3), SUR1 (n=2), GPB2 (n=2), relative to unmutated reference strains. Light gray bars represent the average fitness of each class of mutants in single drug environments, dark gray bars represent fitness predictions in double drug environments made using an additive model, and colored bars represent average fitness in double drug environments (colored blue when fitness is lower than prediction and red when fitness exceeds the prediction). Colors lighten when within 0.5 of the expected value. The type and magnitude of ExE interaction appears to be similar across mutations to the same gene, but different across mutations to different genes. Each row corresponds to one of the double drug environments we study, including **(A)** LRLF, **(B)** LRHF, **(C)** HRLF, **(D)** HRHF. **(E)** ExE for 774 mutants in each studied drug combination broken down by cluster assigned in previous work (56). Mutants are colored by their type of ExE interaction. Here, mutants that experience synergistic interactions are noted with a blue point while antagonistic interactions are noted with a red point. Colors lighten as ExE approaches zero. Sequenced mutants from A-D are shown by colored diamonds. Boxplots summarize the distribution across lower hinges), and highest value within 1.5 × IQR (whiskers).

Overall, we find that mutations to the same gene tend to have similar ExE interactions (**Figure 3A – D**). For example, the 35 PDR1/3 mutants tend to have lower fitness than expected by an additive model in the LRHF environment (**Figure 3A**; left), but not to the same degree as do IRA1 mutants, some of which actually have a slight disadvantage in that double drug environment despite being adaptive in both single drug conditions (**Figure 3A**; middle). And in a different double drug environment, the fitness of all evolved yeast strains with mutations to either PDR1 or PDR 3 is fairly well predicted by an additive model (Figure 3B; left). But an additive model dramatically underestimates the fitness of mutations to the SUR1 gene in the same environment (Figure 3B; right). Across all four double drug environments and all 4 common targets of adaptation we sequenced, the type and magnitude of ExE interactions depends on which gene is mutated (**Figure 3A – D**).

Our observation that ExE varies across mutants does not necessarily arise because we collected adaptive mutants across 12 different selective pressures (56). Mutants that emerge in response to the same selection pressure can have different ExE. For example, IRA1 and GPB2 are both negative regulators of glucose signaling, and both are common targets of adaptation in response to glucose limitation (56, 74, 75). Here, we show that these genes demonstrate different ExE interactions. IRA1 mutants perform worse than expected in LRHF, while GPB2 mutants perform better than expected given their meager fitness advantages in the relevant single drug conditions (**Figure 3B**).

In terms of synergy vs antagonism, our results suggest that a small number of mutations can change a drug combination from having a synergistic to an antagonistic effect. For example, **figure 2C** shows a case where LRLF acts synergistically on a yeast strain harboring a single nucleotide mutation to the HDA1 gene, but acts antagonistically on a different evolved yeast mutant. Similarly, **figure 3** shows cases where a drug pair changes from having a synergistic to an antagonistic effect across different mutants. The extreme sensitivity of synergy to the effect of single mutations has important implications for the development of multidrug strategies that rely on drugs having synergistic or antagonistic effects.

### Some mutants may predict the ExE of other mutants

The above observations highlight the prevalence of ExExG. They beg questions about to what extent there are trends that can help us predict ExE of some mutants from other mutants. These observations also beg questions about the underlying cellular mechanisms that cause ExE interactions to change from one mutant to the next. Both types of questions are related because mutations that affect drug resistance through similar cellular mechanisms may have similar ExE, such that understanding the mechanisms underlying ExE may help predict its direction and magnitude.

We previously showed that many (774) of the yeast strains we study cluster into a small number of groups (6) that each may affect fitness via distinct cellular mechanisms (56). Here, we find that mutants from the same cluster tend to have more similar ExE (**Figure 3E**). For example, the two yeast strains with mutations to SUR1 (Figure 3) clustered together with 107 other strains that have fitness advantages in low (but not high) concentrations of fluconazole (**Figure 3E**; cluster 1) (56). On average, ExE interactions across these 109 yeast strains are predicted by the behavior of the SUR1 strains in figure 3; they tend to behave synergistically in drug combinations containing low flu (**Figure 3E**; cluster 1 in LRLF HRLF), and antagonistically in combinations containing high flu (**Figure 3E**; cluster 1 in LRHF HRHF). Similarly, 31 of the 35 yeast strains with mutations to either PDR1 or PDR3 clustered together with 127 other yeast strains that have fitness advantages in all single and double drug environments (**Figure 3E**; cluster 3) (56). On average, ExE interactions across these strains are predicted by the behavior of the PDR strains in figure 3; they are sometimes synergistic (**Figure 3E**; cluster 3 in HRLF HRHF). This synergism (i.e., mutants are less fit than predicted by an additive model) seems consistent with the mechanism underlying drug resistance in PDR strains. PDR1 and PDR3 regulate a pump that eliminates drugs from cells (76, 77). Perhaps the rate at which this pump removes drug from cells does not increase linearly as more drug is added, therefore an additive model overestimates fitness in double drug environments.

### Considering ExExG suggests a nuanced model for predicting ExE

Modeling ExE in the same way that genetic interactions are modeled may improve predictions. For example, we found it surprising when some mutants that resisted two single drugs lost their fitness advantage when those single drugs were combined (**Figure 4A**; left). However, this loss of fitness is sometimes predictable when we modify GxG (i.e.epistasis) models to study ExE (**Figure 4**; left side). The key is that GxG models incorporate information from a wildtype individual (**Figure 4B**). We can modify this GxG framework to model ExE by incorporating information from an environment lacking drugs. This lets us model the “effect” of each single drug relative to the no drug condition similarly to how models of GxG model the “effect” of each single mutation relative to the wildtype (12) (**Figure 4B – C**). Once this effect is measured, it creates an expectation for how addition of this drug will modify fitness (**Figure 4C**; purple diamond). We call our model the “Drug Effect” (DE) model because, like the GxG framework upon which it is based, it assumes that a perturbation (e.g., a drug) has a static effect on a given mutant’s fitness. One limitation however, is that to implement this DE model, one must have fitness measurements not only from single drug and double drug conditions, but also in conditions lacking any drug.

**Figure 4:**
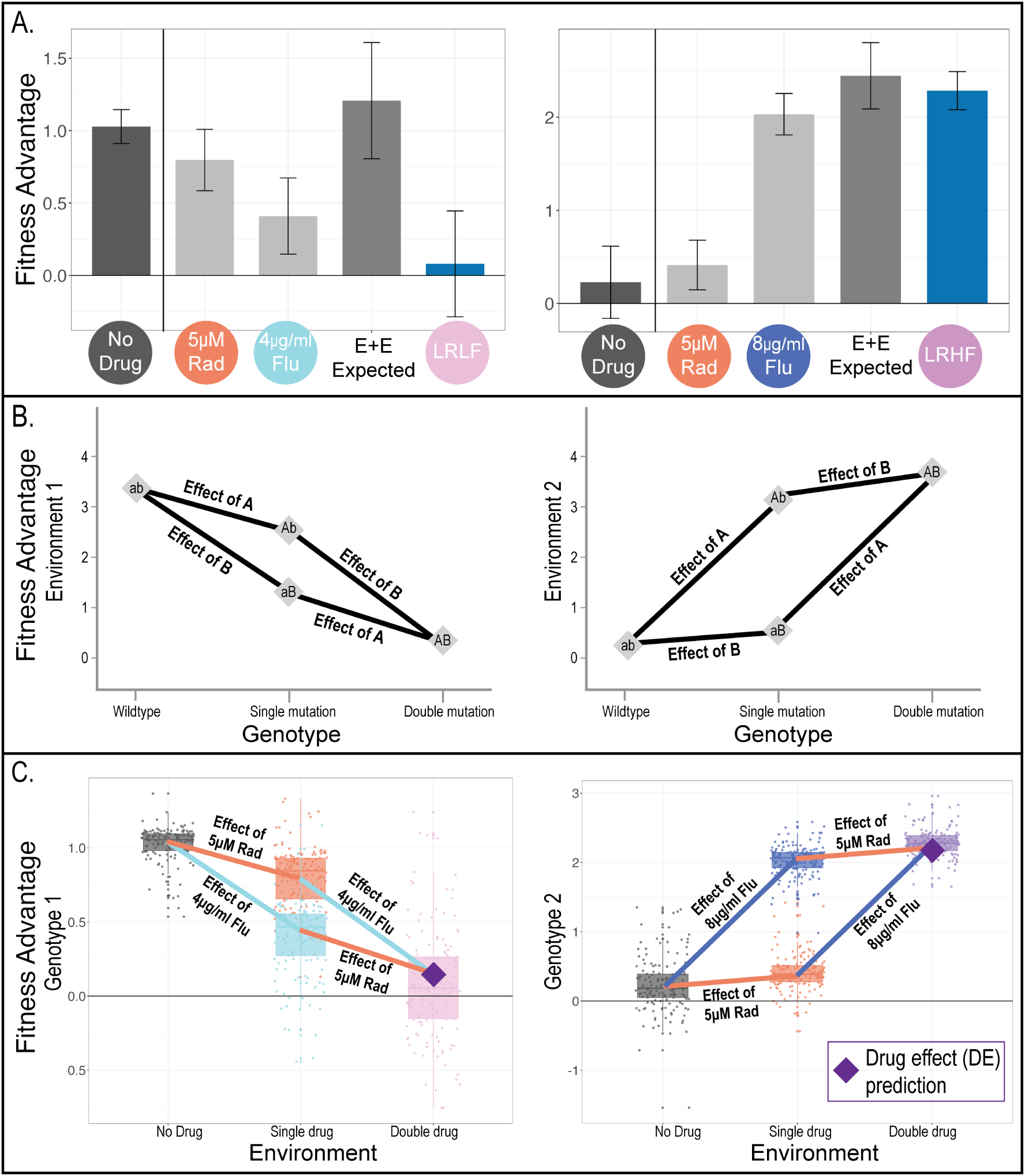
Classical GxG framework inspired a new “drug effect” (DE) model that accurately predicts the behavior of some drug resistant mutants in double drug environments. **(A)** Our original additive model (“E+E”) makes poor fitness predictions for the 145 mutants in the left panel, but not for the 158 mutants in the right panel. Another key difference is that the mutants in the left panel have fitness advantages over the reference strain in the no drug environment, while the mutants in the right panel do not. The mutants in each panel clustered together in previous work based on their fitness in 12 environments (56). Dark gray bars represent average fitness in no drug, light gray bars represent average fitness in single drug environments, medium gray bars represent fitness predictions in double drug environments made using our original additive model, and colored bars represent average fitness in double drug environments. Error bars represent standard deviation. **(B)** Classic GxG additive models are different from the additive models in **panel A** and in earlier figures. GxG models add together the effect of each single mutation to predict the fitness of the double mutant, rather than adding together the fitness of each single mutant (12). The left panel provides an example where the wildtype (ab) has a fitness advantage in environment 1. Gaining mutation A or B results in decreased fitness. Subtracting the effect of both A and B allows for the correct prediction of the double mutant’s (AB) fitness in environment 1. The right panel presents a second environment where the wildtype fitness is improved by mutations A and B. Here adding the effect of both A and B results in accurate prediction of the double mutant’s fitness. **(C)** Repurposing the GxG model in **panel B** to predict fitness results in accurate predictions for the mutants described in panel A. Boxplots summarize the distribution across all mutants, displaying the median (center line), interquartile range (IQR) (upper and lower hinges), and highest value within 1.5 × IQR (whiskers). No drug is shown in dark gray, single drugs in blue/orange and double drugs in pink/purple. The effect of each drug is represented by a colored line matching that of the single drug. The average prediction of the DE model for both groups of mutants is shown by a purple diamond.

To better illustrate the DE model, consider that the decisive difference between the mutants in **figure 4A** left and right is their fitness in conditions lacking any drug. The mutants on the left have a fitness advantage in conditions lacking drug (**Figure 4A**; no drug). While the mutants on the left also have a fitness advantage in each single drug, the “effect” of each single drug on fitness is actually negative. In other words, these drugs reduce the fitness advantage. The DE model thus correctly predicts that the effect of combining both drugs will be a further reduction in fitness (**Figure 4C**; left) while our original additive model fails to make an accurate prediction (**Figure 4A**; left). But the mutants on the right have no advantage in the no drug environment, and the “effect” of adding each single drug is actually to improve their relative fitness (**Figure 4A**; right). Here, the DE model performs similarly to our original additive model in predicting fitness in the multidrug environment (**Figure 4**; right). An important caveat is that, although the DE framework makes reasonable fitness predictions for these two drug pairs, it fails in many other environments and for many other genotypes, again highlighting the prevalence of ExExG (**Figure S1**).

### Different mutants have different drug dose-response curves

One major model for predicting ExE that we do not utilize in **figure 2B** (or elsewhere) is Loewe additivity. This model allows for non-linear dose-response curves when predicting how environments interact. Consider the simplest case where the two environments in question are actually two different concentrations of the same drug. The effect of combining these environments might not be predicted by an additive model if the response curve to this drug is non-linear (**Figure 5A**). Just as nonlinearities can lead to the appearance of ExE, they also commonly result in GxG (**Figure 5B**) (12).

**Figure 5:**
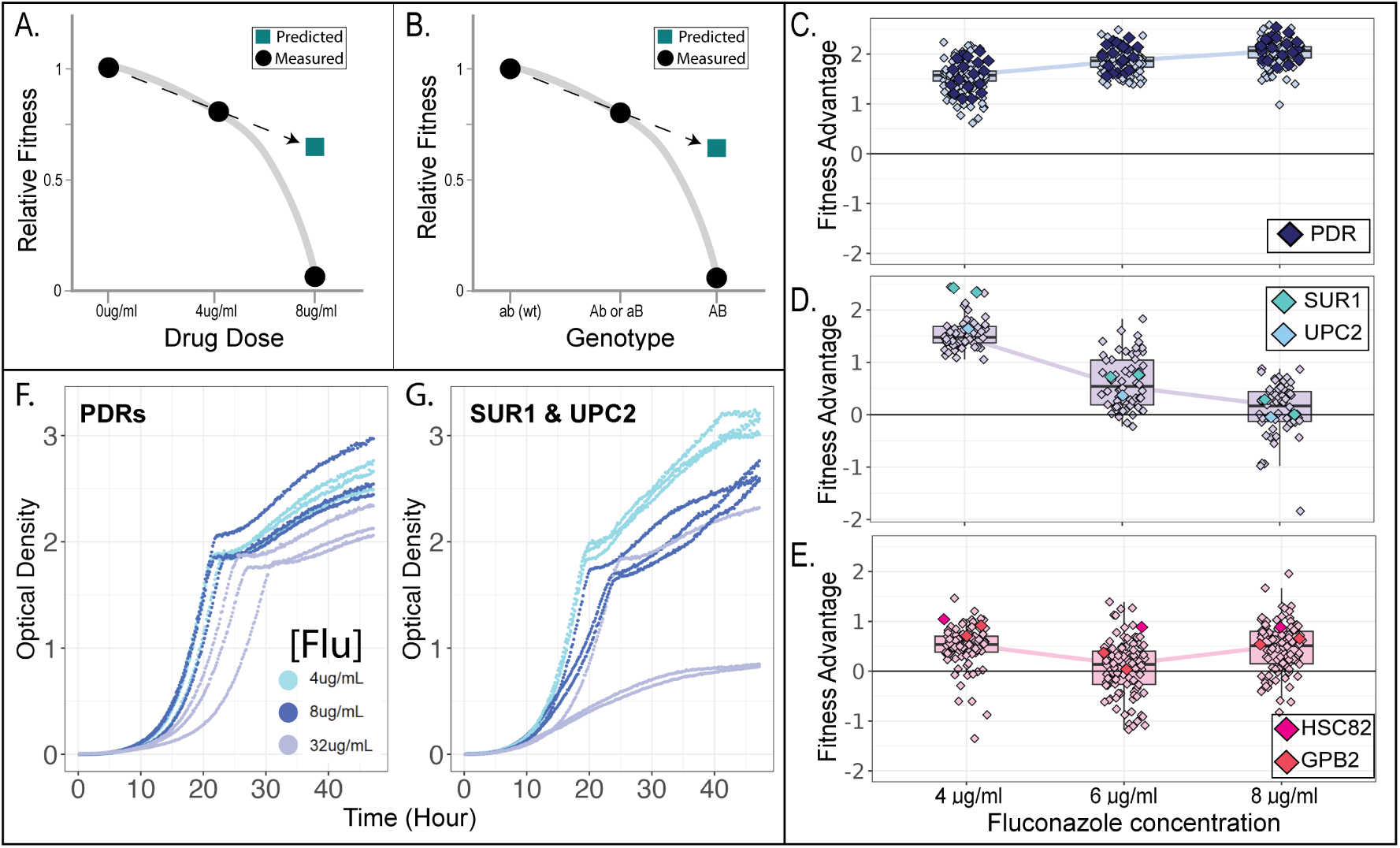
Different drug-resistant mutants have different drug dose responses. (A–B) Toy examples showing how fitness predictions made assuming an additive model can fail when nonlinearities are present. **(C–E)** A simple nonlinear model cannot account for ExE in these data because different mutants have different drug dose responses. Each panel captures unique mutants; sequenced mutants are highlighted with diamonds corresponding in color to those in figure 3. Boxplots summarize each distribution, displaying the median (center line), interquartile range (IQR) (upper and lower hinges), and highest value within 1.5 × IQR (whiskers). **(F)** Three isolated mutants from **panel C** have similar growth curves in multiple fluconazole concentrations. **(G)** Three isolated mutants from **panel D** grow better in low fluconazole and increasingly worse as the drug concentration increases.

We cannot use Loewe additivity to capture nonlinearities because some of our mutants have extremely different dose-response curves than others. For example, we see a distinct class of mutants for which relative fitness increases with the concentration of fluconazole (**Figure 5C**), another class for which fitness decreases with the concentration of fluconazole (**Figure 5D**), and still another class for which fitness is similar in low and high fluconazole conditions (**Figure 5E**). No single non-linear dose-response curve can describe how fitness changes upon combining two different concentrations of fluconazole for all of these mutants. Instead, we again conclude that multiple different models of how environments interact are required to capture the behavior of these diverse mutants (in other words, we conclude that there is ExExG).

One question that may arise is: to what extent does our decision to study relative fitness advantages affect our results. Many previous studies on ExE and GxG interactions focus on relative fitness (13, 32, 33, 78), e.g. by measuring growth relative to a condition without drugs (Figure 5A) and growth relative to a strain without mutations (**Figure 5B**). In our study, we did not measure growth curves for each mutant, but instead conducted pooled fitness competitions (56), calculating fitness advantages of all mutants relative to an unmutated reference strain, and comparing these advantages across environments. This can make it harder to interpret the drugdose response curves we see in **figures 5C – E**. For example, the increase in relative fitness advantage across conditions observed in **figure 5C** may indicate that these mutants perform better as the drug concentration increases. Alternatively, their growth rate might be insensitive to changes in the drug concentration, and their increased fitness advantage could reflect the worsening performance of the unmutated reference strain. Indeed, we find that the latter is true. When we previously measured the growth rates of three isolates each with a mutation to either the PDR1 or PDR3 genes, we find that these mutants have similar growth curves in a range of fluconazole concentrations (Figure 5F) (56). On the other hand, the three isolated mutants depicted in Figure 5D with mutations to either SUR1 or UPC2 perform more poorly as fluconazole concentrations increase (**Figure 5G**) (56). Whether models seeking to predict how microorganisms will respond to drug treatment should focus on absolute measures of performance, such as the growth rate of isolated cultures, or relative measures of fitness, such as the advantage in a pooled competition, is a question for another study, though both seem very important (56, 79–81). The salient point, with respect to this study, is that these two groups of mutants behave differently, in both their absolute (**Figure 5F–G**) and relative fitness (**Figure 5C– D**), in their responses to increasing fluconazole concentrations (**Figure 5**) and their responses to multidrug environments (**Figure 3**), signifying the presence of ExExG.

## Discussion

In this study, we explored ExE interactions (i.e. drug interactions) in a large population of drug resistant yeast strains and found that different strains often have different ExE, meaning that ExExG is common. In other words, the way two drugs interact, whether their combined effect is stronger or weaker than the sum of their individual effects, depends on genotype. This means that we may require multiple different models to predict the way the fitness of a collection of mutants will respond to combined drug treatment. For example, three different models are needed to predict how the fitness of three different groups of mutants responds to increased fluconazole concentrations (**Figure 5 C – E**). And our DE model predicts the fitness decrease observed in multidrug conditions that was unexpected under a more simplistic additive model (**Figure 4**), but does so only for some mutants (Figure S1). There are hints of predictability in that some drugresistant yeast strains, such as those with mutations to the same genes, tend to have similar ExE interactions (**Figure 3**). In sum, this work suggests that in order to make better predictions about ExE interactions, including

### Is it useful to create a new term, “ExExG”?

When building predictive models of interactions, it may be helpful to consider when it is useful to codify contextual perturbations as genetic vs. environmental or otherwise? On one hand, classifying which studies focus on GxG, GxE, GxGxE, ExExG, etc, is tedious and can be confusing (but we hope Figure 1 will help). Further, classifying based on these factors can create a language barrier whereby studies focusing on drug interactions are disparate from those focusing on genetic interactions. Here we show that communication between these fields is important by demonstrating that classical models of genetic interactions can be helpful in understanding drug interactions (**Figure 4**). Finally, genetic and environmental perturbations are indeed similar in that they can both change the way genotype maps to phenotype, therefore, perhaps they should be modeled in the same ways simply as “perturbations” that affect phenotype, or as “parcels of information” that are interpreted by cells and manifest in phenotypic outcomes (11). On the other hand, when asking more specific questions pertaining to specific genetic or environmental factors, distinguishing contexts is important.

### Why study ExE and ExExG?

A key reason to study ExE (or other) interactions is a desire to identify rules operating in biological systems that allow for better predictions of their behavior (e.g., phenotype) based on different factors. For example, if we knew that two drugs interact synergistically, we could predict that together they would be more effective for treating infections. Several modern paradigms aim to add rhyme and reason to even nonlinear interactions. One perspective, labeled “global” or “nonspecific” epistasis, posits that the even non-additive interactions between perturbations or parcels can follow a mathematical pattern, which offers hope that we might one day truly predict how systems work (12, 82–84).

High throughput technologies that survey genotype and phenotype with increasingly fine levels of detail could help resolve the complexity and caprice of biological systems in the form of basic rules. But in biology and other disciplines, we know that rules often do not apply to every circumstance. One might even suggest that biology has become a field defined by an understanding of the context-dependence of its basic axioms (5). In this study, we find that rules governing how drugs interact (and models based upon those rules) do not apply to all mutants. If this departure from the convention were isolated to a small group of mutants, then perhaps elucidating general rules would still be possible or useful. But if each mutant needs its own rule to describe ExE interactions, then the generality of these principles can be called into question. On the other hand, even in cases where interactions undermine neat predictions, some previous work suggests that not all aspects of a system must be well known or behaved in order to develop a reasonably predictive set of rules (31, 57, 62, 74, 85). Our study suggests that more work is needed to understand the complexity of biological systems (56, 74, 86) and the extent to which rules can generate predictions that capture their behavior.

## Methods

### Data acquired from experimental evolutions and fitness competitions

All data presented in this work was collected as previously described in (Schmidlin et al., 2024). Briefly, 300,000 barcoded yeast lineages were evolved for 7 weeks in 10 drug conditions and 2 controls. From these evolutions, 21,000 (2k from each evolution) colonies were selected for a fitness remeasurement experiment. Barcode sequencing was performed every 48 hours and log-linear changes in barcode frequencies over 4 time points were used to infer fitness. From this subset, a final collection of 774 lineages, characterized by greater than 500 barcode reads from each of the 12 environments, were analyzed from this previous study. However, there are additional lineages that have greater than 500 barcode reads/condition if you require fewer conditions. Since we were interested in ExE interactions, we created four improved datasets that contained lineages present in the no drug control, both single drugs that made up the combination and the double drug combination. Datasets were improved as follows: LRLF: n=1688; LRHF: n=850; HRLF: n=1318; HRHF: n=1023.

### Definitions for drug interaction models

Several models were used to quantify drug interactions and are defined as follows:

1 Additive Model (E+E): The fitness of each lineage in the defined drug combination is determined by the sum of the relative fitness values in drug environment 1 and drug environment 2. For our work here, this constitutes the expected model.
2 Bliss Independence Model (Bliss): Prior to calculation, each fitness value was converted to a percentage based on the maximum observed fitness value in the respective drug combination (DC). The formula is as follows: (Fitness in drug environment 1 + fitness in drug environment 2 (Fitness in drug environment 1* fitness in drug environment 2))*maxDC.
3 Highest Single Agent Model (HSA): This model reports the maximum fitness value among the single drugs present in the combination.
4 Average Model (Avg): The model fitness in the drug combination is represented as an average between the two single drugs.
5 Drug Effect Model (DE): This model first finds the fitness value for a single drug, then from this value subtracts the fitness of the lineage in no drug from the fitness of the lineage in the second single drug. The result is the prediction for the drug combination.

All code is available on OSF under the project: Environment by environment interactions (ExE) differ across genetic backgrounds (ExExG).

### Quantifying ExE for 774 lineages in four drug combinations

In order to quantify the amount of ExE captured in our dataset, we first estimated the fitness of each lineage in the four drug combination environments using log linear slope as previously described (Schmidlin et al., 2024). Five predictions, one for each model above, were made for each lineage in the dataset. Once predictions were calculated, they were subtracted from the known fitness. Differences that did not equal 0 (truth minus prediction) were considered to have environment by environment interactions and are reported as ExE.

## SUPPLEMENT

**Table 1.**
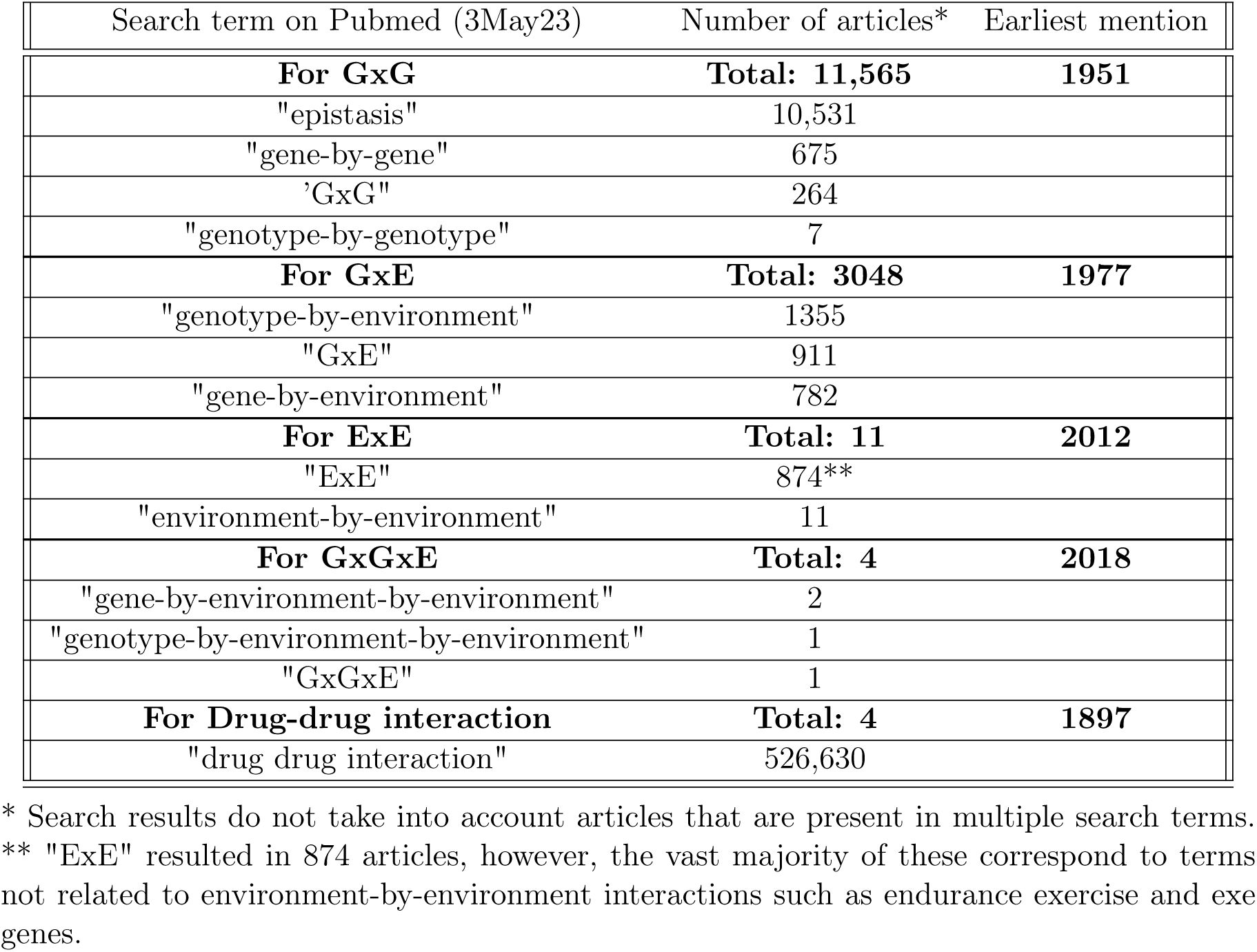
Pubmed search terms and results for Figure 1A.

| Search term on Pubmed (3May23) | Number of articles* | Earliest mention |
| --- | --- | --- |
| **For GxG** | **Total: 11,565** | **1951** |
| "epistasis" | 10,531 |  |
| "gene-by-gene" | 675 |  |
| 'GxG' | 264 |  |
| "genotype-by-genotype" | 7 |  |
| **For GxE** | **Total: 3048** | **1977** |
| "genotype-by-environment" | 1355 |  |
| "GxE" | 911 |  |
| "gene-by-environment" | 782 |  |
| **For ExE** | **Total: 11** | **2012** |
| "ExE" | 874** |  |
| "environment-by-environment" | 11 |  |
| **For GxGxE** | **Total: 4** | **2018** |
| "gene-by-environment-by-environment" | 2 |  |
| "genotype-by-environment-by-environment" | 1 |  |
| "GxGxE" | 1 |  |
| **For Drug-drug interaction** | **Total: 4** | **1897** |
| "drug drug interaction" | 526,630 |  |

\* Search results do not take into account articles that are present in multiple search terms.
\*\* "ExE" resulted in 874 articles, however, the vast majority of these correspond to terms not related to environment-by-environment interactions such as endurance exercise and exe genes.

**Figure S1:**
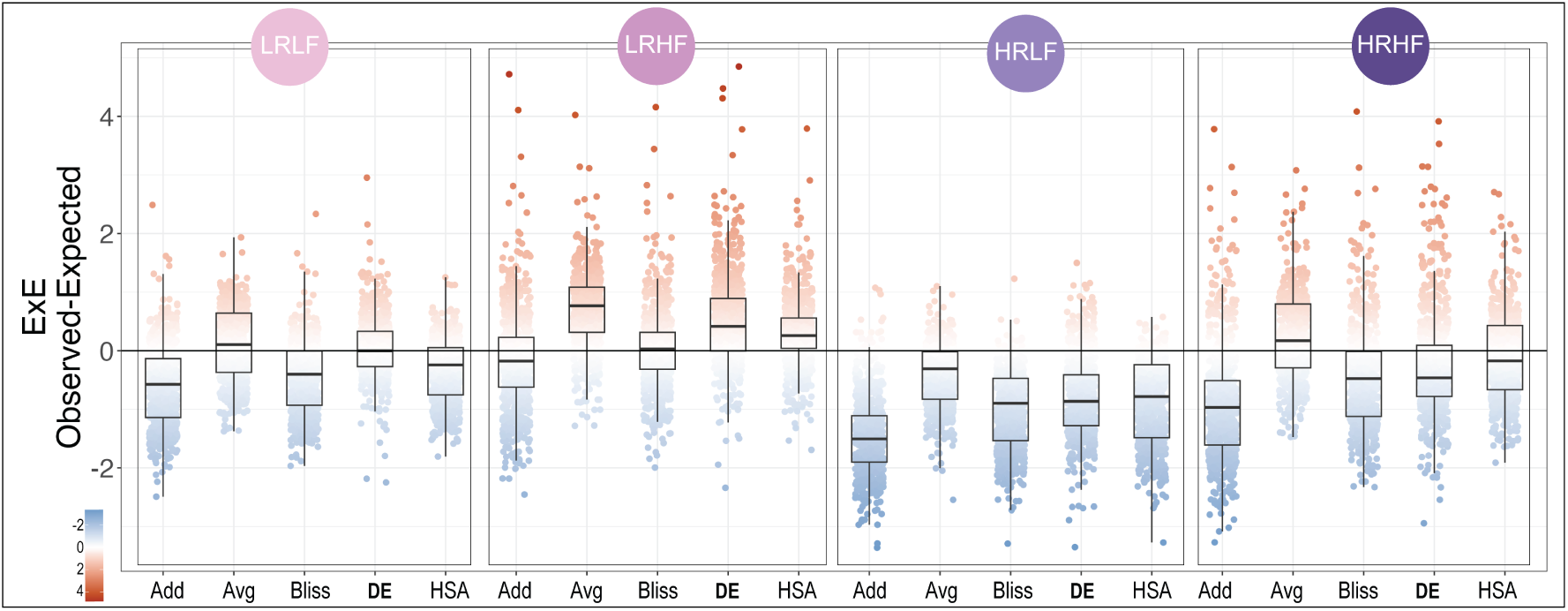
The DE model predicts ExE for some mutants but not others.

